# Thermoneutral Housing has Limited Effects on Social Isolation-Induced Bone Loss in Male C57BL/6J Mice

**DOI:** 10.1101/2024.08.09.607315

**Authors:** Rebecca V. Mountain, Rebecca L. Peters, Audrie L. Langlais, J. Patrizia Stohn, Christine W. Lary, Katherine J. Motyl

## Abstract

Social isolation stress has numerous known negative health effects, including increased risk for cardiovascular disease, dementia, as well as overall mortality. The impacts of social isolation on skeletal health, however, have not been thoroughly investigated. We previously found that four weeks of social isolation through single housing led to a significant reduction in trabecular and cortical bone in male, but not female, mice. One possible explanation for these changes in male mice is thermal stress due to sub-thermoneutral housing. Single housing at room temperature (∼20-25°C)—below the thermoneutral range of mice (∼26-34°C)—may lead to cold stress, which has known negative effects on bone. Therefore, the aim of this study was to test the hypothesis that housing mice near thermoneutrality, thereby ameliorating cold-stress, will prevent social isolation-induced bone loss in male C57BL/6J mice. 16-week-old mice were randomized into social isolation (1 mouse/cage) or grouped housing (4 mice/cage) at either room temperature (∼23°C) or in a warm temperature incubator (∼28°C) for four weeks (N=8/group). As seen in our previous studies, isolated mice at room temperature had significantly reduced bone parameters, including femoral bone volume fraction (BV/TV), bone mineral density (BMD), and cortical thickness. Contrary to our hypothesis, these negative effects on bone were not ameliorated by thermoneutral housing. Social isolation increased glucocorticoid-related gene expression in bone and *Ucp1* and *Pdk4* expression in BAT across temperatures, while thermoneutral housing increased percent lipid area and decreased *Ucp1* and *Pdk4* expression in BAT across housing conditions. Overall, our data suggest social isolation-induced bone loss is not a result of thermal stress from single housing and provides a key insight into the mechanism mediating the effects of isolation on skeletal health.

**Lay Summary:** Social isolation is a major public health concern and is known to increase the risk for many diseases, including heart disease and dementia. The impact of social isolation on bone health, however, has not been well-studied. We previously found that four weeks of social isolation reduces bone in male mice. Isolated mice may experience more cold stress than mice housed in groups, as we commonly keep laboratory mice at temperatures below their ideal range, which could lead to bone loss. The aim of our study was therefore to test if housing mice at warmer temperatures, within their ideal temperature range, prevents isolation-induced bone loss in male mice. We found that housing mice at warmer temperatures did not fully prevent isolation-induced bone loss. We also found social isolation increased the expression of genes related to glucocorticoid signaling in bone across temperatures, as well as genes associated with mitochondrial metabolism within fat tissue. Overall, our results show that social isolation-induced bone loss is likely not a result of cold stress from single housing and provide insight into the mechanisms by which isolation causes bone loss.

## 1. Introduction

Social isolation and loneliness, as well as their associated health consequences, have gained increased attention from both public health officials and the mass media as a result of the global COVID-19 pandemic. While the increase in isolation and loneliness had been a concern for many years, affecting 1 in 4 adults over the age of 65^(1)^, the pandemic exacerbated rates of isolation^(2)^ and concerns about the public health impact across age demographics. The United States surgeon general recently released a report in 2023 highlighting the dangers of social isolation and importance of social connection for public health^(3)^. Social isolation is associated with a significant increase in cardiovascular disease risk^(4)^, neurodegenerative disease^(5,6)^, and mental health disorders^(7)^. Isolation is also associated with a substantial increase in mortality risk, exceeding that of chronic smoking or drinking^(8)^.

While there has been an abundance of research on the effects of isolation on other physiological systems, including the cardiovascular system, there has been very little research on the effects of isolation on skeletal health^(9,10)^. We previously tested the effects of four weeks of social isolation through single housing in 16-week-old male and female C57BL/6J mice. We found that four weeks of social isolation led to a significant reduction in trabecular (26%) and cortical (9%) bone in male, but not female, mice^(11)^. We further found evidence of reduced bone turnover in the male mice, with reduced formation (e.g., *Runx2*, *Dmp1*) and resorption-related (e.g., *Ctsk, Acp5*) gene expression, as well as a trending decrease in osteoblast number and surface area.

One possible explanation for the negative effects of isolation on bone in male mice is the potential for thermal stress during single housing. Mice are typically housed in research facilities at room temperature (∼20-25°C) which is below the thermoneutral range of mice (∼26-34°C), or the temperature at which they maintain their body temperature without increasing their metabolic rate^(12)^. Previous studies have demonstrated that thermal stress as a result of sub-thermoneutral housing can result in premature bone loss, as well as increased brown adipose tissue (BAT) activation^(13,14)^. Singly housed mice may be particularly vulnerable to this thermal stress as they lack the additional body heat of cage mates, which could exacerbate bone loss.

The aim of this study was to investigate the role of housing temperature on social isolation-induced bone loss. We tested the hypothesis that housing mice at thermoneutrality, thereby ameliorating cold-stress, will prevent social isolation-induced bone loss in male C57BL/6J mice. This is the first study to investigate the interaction of housing temperature and isolation on bone metabolism. The results of this project will aid in identifying the mechanisms involved in isolation-induced bone loss, potentially leading to future interventions or therapeutic targets.

## 2. Materials and Methods

### 2.1 Mice

10-week-old male C57BL/6J mice (N=32) were obtained from the Jackson Laboratory (Strain #000664, Bar Harbor, ME), and acclimated for six weeks after arrival to eliminate any potential stress-related effects from shipment. All mice were housed in the barrier animal facility at MaineHealth Institute for Research (MHIR). MHIR is an Association for Assessment and Accreditation of Laboratory Animal Care (AAALAC) accredited facility, and all procedures utilized in this study were approved by the MHIR Institutional Animal Care and Use Committee (IACUC). All mice were kept on a 14 hr light/10 hr dark cycle and provided regular chow (Teklad global 18% protein diet, #2918, Envigo, Indianapolis, IN, USA) and water *ad libitum*.

### 2.2 Experimental Design

Mice were grouped or isolated as described in our previous publication^(11)^. Briefly, at 16 weeks of age, mice were randomized into either control/grouped (4 mice/cage) or isolated (1 mouse/cage) housing for four weeks with a shepherd shack in each cage for enrichment for both groups. Half of each housing group (grouped or isolated) (N=8/group) were housed at room temperature (∼23°C) in a standard room within our barrier facility. The other half were housed in a warm temperature incubator (Model RIS52SD, Powers Scientific Inc.) set for 29 ± 1° C with a minimum 30% humidity^(15)^. The precise thermoneutral range can vary by mouse strain and age among other factors, however, we selected 29 ± 1°C based on previous literature rather than performing colorimetric or temperature preference tests. While recognizing this limitation, we refer to this temperature as “thermoneutral” or “thermoneutrality” in the remainder of this paper for simplicity. Temperature and humidity were confirmed on both digital and manual gauges twice daily and averaged ∼28°C and ∼34% humidity over the course of the four weeks. Both room temperature and incubator mice were kept on the same light/dark cycle and food as mentioned above. Cage changes were performed once a week for all groups of mice.

### 2.3 Dual-energy x-ray absorptiometry (DXA)

Dual energy x-ray absorptiometry (DXA) was performed at baseline (16 weeks of age) and endpoint (20 weeks of age), using a Hologic Faxitron UltraFocus DXA system. Bone and fat phantoms provided by the manufacturer were used to calibrate the system before each scanning session. Mice were weighed, and areal total body (post-cranial), femoral, and vertebral (fifth lumbar) bone mineral density (aBMD) were measured as well as total body lean and fat mass. Baseline DXA was performed to ensure there were no significant mean differences in bone or body parameters between experimental groups before the experiment began.

### 2.4 Micro-computed Tomography (μCT)

Trabecular and cortical bone volume, mineral density, and microarchitecture were measured using high resolution microcomputed tomography (μCT) with a resolution of 10.5 μm (vivaCT 40, Scanco Medical AG, Brüttisellen, Switzerland). Femurs and the vertebral column were dissected after sacrifice, fixed in formalin (10% Neutral buffered) for 48 hours, and then transferred to 70% ethanol. Cortical bone was assessed at the femoral mid-diaphysis, and trabecular architecture was assessed at the distal femur metaphysis and the L5 vertebral body. All bone scans were acquired with an isotropic voxel size of 10.5 μm^3^, 70 kVp peak x-ray tube intensity, a 114 mA x-ray tube current, and 250 ms integration time. For each scan four femora were evenly distributed in a custom-made poly ether imide sample holder, while the vertebral columns were scanned individually. Gassuian filtration and segmentation was performed on all scans, and all analyses were performed using Scanco µCT Evaluation ProgramV6.6.

### 2.5 Bone Turnover Markers

After decapitation, blood was collected from all mice and allowed to clot at room temperature for at least 10 minutes. Blood was centrifuged at 10,000 rpm for 10 minutes, and serum was isolated and stored at -80°C. Serum CTX-I and P1NP concentration were measured with the RatLaps CTX-I and Rat/Mouse P1NP enzyme immunoassays (EIA, Immunodiagnostic Systems, United Kingdom) respectively. Assays were performed following the manufacturer’s instructions in duplicate and read on a FlexStation 3 plate reader (Molecular Devices, Eugene, OR, USA). Results were determined using the FlexStation software with a 4-parameter logistic curve. Samples with a coefficient of variation exceeding 20% were excluded from analysis resulting in a N=4-8 per group.

### 2.6 Brown Adipose Tissue Histology

After dissection, brown adipose tissue was fixed in formalin (10%) for 48 hours and transferred to 70% dehydrant alcohol. Samples were then embedded in paraffin and cut to 5µm sections and stained with hematoxylin and eosin (H&E) by the MHIR Histology Core following standard protocols (N=8/group). Representative images were taken with the Keyence BZ-X800 series Microscope in the MHIR Confocal Microscopy Core at 10x magnification. Images were then quantified in FIJI following the protocol by Tero et al 2022^(16)^. In brief, images were segmented, non-tissue segments were converted to black, and an Otsu threshold was applied to allow for the differentiation of lipid space versus the total area.

### 2.7 RNA Isolation and Real-time qPCR (RT-qPCR)

Whole tibia and brown adipose tissue were collected, and flash frozen in liquid nitrogen and stored at -80°C (N=8/group). Samples were crushed under liquid nitrogen conditions and homogenized in 1mL of TriReagent (MRC, Cincinnati, OH). Samples were incubated in 200 µL of chloroform for 10 minutes at room temperature, and then centrifuged at 12,000 rpm for 15 minutes at 4°C. The aqueous layer was isolated and mixed with 500 µL of isopropanol. Samples were then frozen overnight at -80°C. The following day, samples were centrifuged at 12,000 rpm for 15 minutes at 4°C, supernatant removed, and the RNA pellet washed in 1 mL of 75% ethanol twice. After removing the supernatant a final time, the pellet was allowed to air dry for 5 minutes and then dissolved in 80 µL or 40 µL of nuclease-free H_2_O for whole bone or brown adipose tissue respectively. Samples were frozen overnight at -80°C. RNA concentration was determined using the NanoDrop 2000 (Thermo Fisher Scientific, Waltham, MA, USA), and diluted if necessary (if >1000ng/µL) with nuclease-free H_2_O. 1000 ng of RNA was added to each cDNA reaction with 10x RT buffer, 25x dNTP, 10x random primers, Reverse Transcriptase, and nuclease-free H_2_O (High Capacity cDNA Reverse Transcription Kit, Thermo Fisher Scientific, Waltham, MA). Samples were then run on a PCR protocol of 10 min at 25°C, 120 min at 37°C, 5 min at 85°C, and cooled to 4°C, and diluted with 180 μL of nuclease-free H_2_O. RT-qPCR was performed using 3 µL cDNA, nuclease-free water, SYBR green, (BioRad, Hercules, CA), appropriate forward and reverse primers, and run on a BioRad Laboratories CFX 384 real-time PCR system. Primers were purchased from Integrated DNA Technologies (Coravile, IA) or Qiagen (Germantown, MD). All primer sequences used in these analyses are listed in Supplementary **Table S1**. Beta actin (*Actb*) was used as the housekeeping gene for whole bone. In the brown adipose tissue, all potential housekeeping genes tested were significantly different between treatment groups. To account for this, and as all cDNA samples had the same quantity of input RNA, gene expression in the brown adipose tissue was normalized to the highest Cq, or lowest expressor, within each gene. Any values greater than 2 standard deviations above or below the group mean were excluded as outliers.

### 2.8 Statistical Analysis

Statistical analyses were performed using GraphPad Prism 10 XML Project® software. Student’s t-test was used to test for significant differences in baseline DXA parameters between experimental groups. Two-way ANOVA was used for all experimental analyses testing the effects of isolation or housing temperature. α ≤ 0.05 was considered statistically significant. Tukey’s post hoc test was performed for multiple comparisons for any data with a significant interaction effect.

## 3. Results

### 3.1 Social isolation reduced trabecular and cortical bone parameters across housing temperatures

Social isolation reduced femoral trabecular and cortical bone parameters across housing temperatures (**Figure 1**; **Tables 1 and 2**). Housing had a significant main effect on femoral BV/TV, BMD, BS/BV, Tb.N. and Tb.Sp. There was also a significant interaction effect for several trabecular parameters (see **Table 1**). Isolation reduced BV/TV and BMD by approximately 35% and 27% respectively at room temperature, but only by 8% for both parameters at thermoneutrality compared to grouped mice. This amelioration of isolation-induced bone loss, however, is driven in major part by a trending reduction in parameters within the grouped-thermoneutral mice relative to the grouped-room temperature mice. While thermoneutrality increased BV/TV and BMD by 20% within the isolated mice, it also *decreased* BV/TV and BMD by 16% and 11% respectively within the grouped-housed mice.

**Figure 1.**
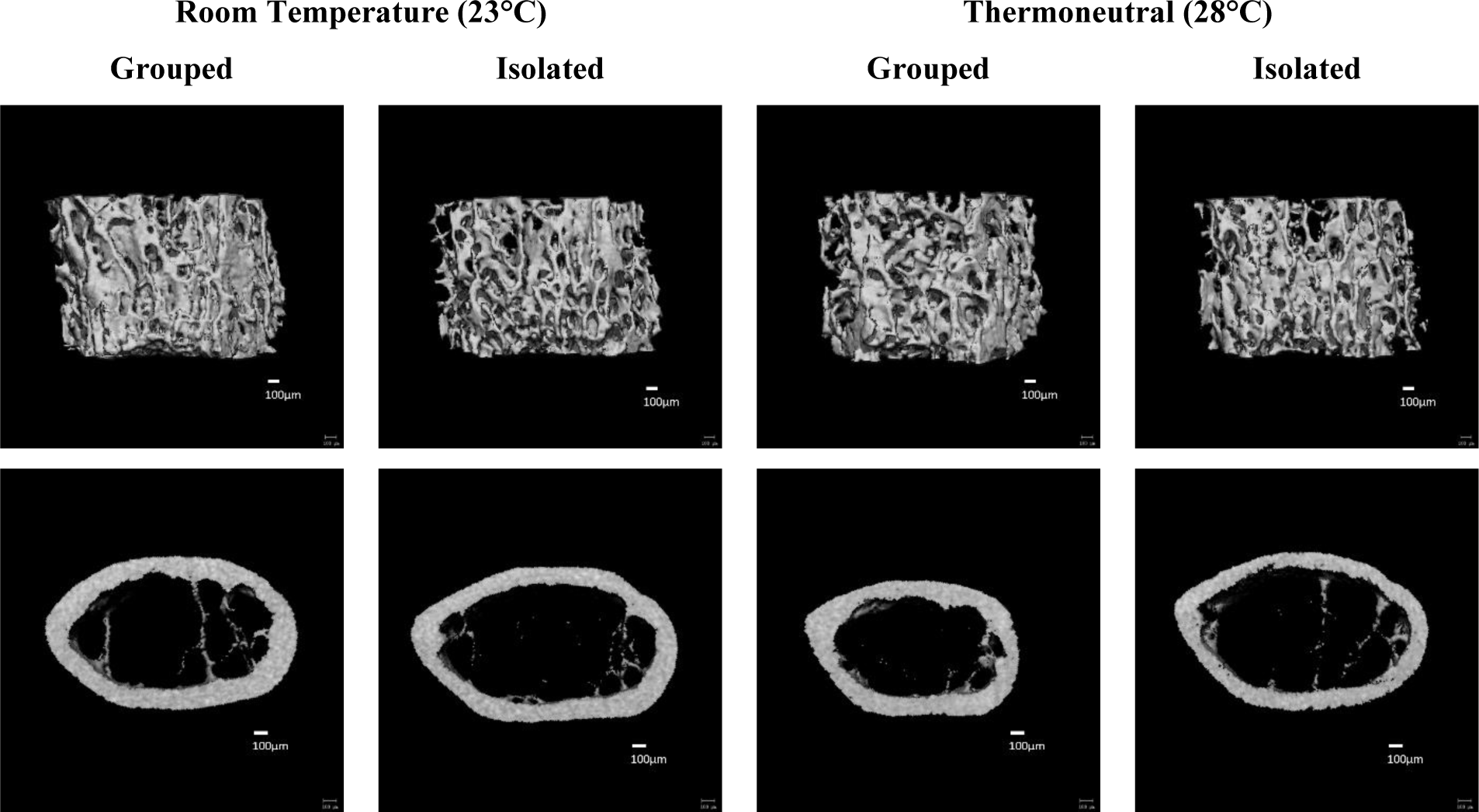
Thermoneutral housing has limited effects on social isolation-induced trabecular bone loss and did not improve cortical bone parameters in isolated mice. 16-week-old male mice were housed at either room temperature or thermoneutral in social isolation (1 mouse/cage) or grouped housing (4 mice/cage) for four weeks. Changes in distal femur trabecular and cortical midshaft bone density and microarchitecture were measured using μCT. Representative images shown. Scale bar = 100μm.

**Table 1.**
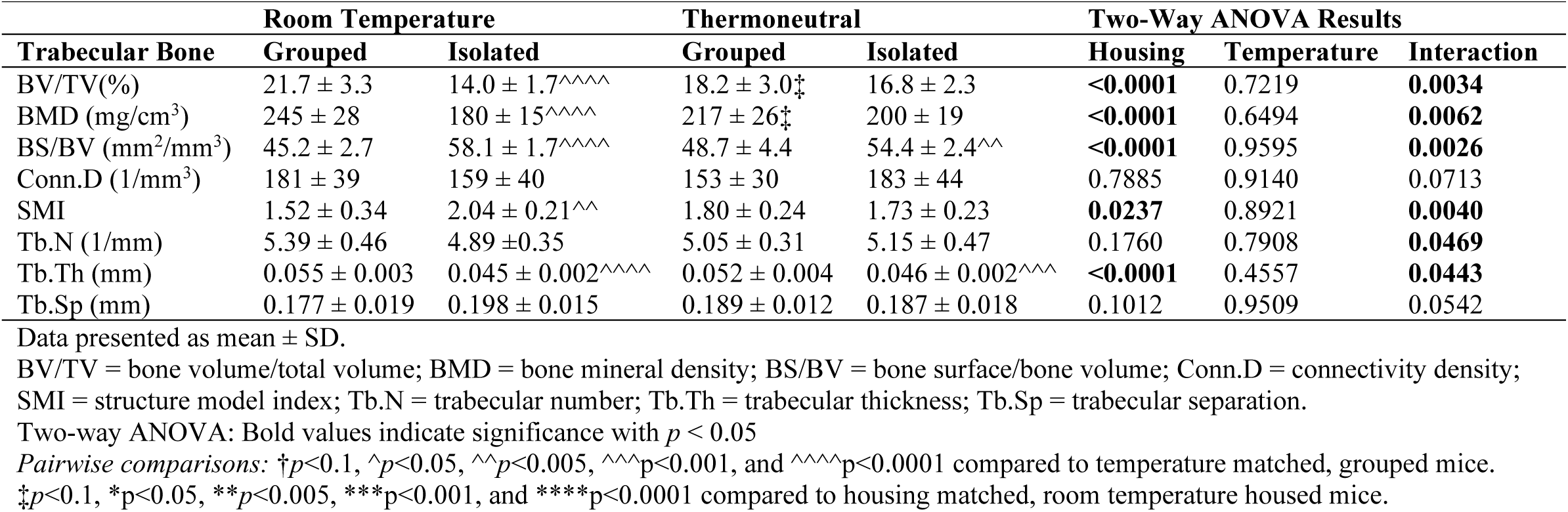
Trabecular microarchitecture of distal femur of male mice housed at either room temperature or thermoneutral in grouped or isolated housing for four weeks. (N=7-8/group).

**Table 2.** Cortical microarchitecture of distal femur of male mice housed at either room temperature or thermoneutral in grouped or isolated housing for four weeks. (N=7-8/group).

| Cortical Bone | Room Temperature |  | Thermoneutral |  | Two-Way ANOVA Results |  |  |
| --- | --- | --- | --- | --- | --- | --- | --- |
|  | Grouped | Isolated | Grouped | Isolated | Housing | Temperature | Interaction |
| Ct.Ar (mm<sup>2</sup>) | 0.937 ± 0.065 | 0.848 ± 0.054 | 0.894 ± 0.058 | 0.856 ± 0.038 | **0.0030** | 0.3840 | 0.2087 |
| Ma.Ar (mm<sup>2</sup>) | 1.12 ± 0.11 | 1.23 ± 0.13 | 1.14 ± 0.15 | 1.30 ± 0.09 | **0.0055** | 0.3026 | 0.5040 |
| Tt.Ar (mm<sup>2</sup>) | 2.06 ± 0.15 | 2.08 ± 0.18 | 2.04 ± 0.19 | 2.16 ± 0.10 | 0.2339 | 0.6151 | 0.3450 |
| Ct.Ar/Tt.Ar (%) | 45.5 ± 2.4 | 40.9 ± 1.2 | 44.1 ± 2.7 | 39.7 ± 2.0 | <**0.0001** | 0.0974 | 0.9205 |
| Ct.Th (mm) | 0.190 ± 0.009 | 0.168 ± 0.006 | 0.180 ± 0.006 | 0.165 ± 0.008 | <**0.0001** | **0.0177** | 0.2095 |
| Ct. TMD (mg/cm<sup>3</sup>) | 1266 ± 11 | 1249 ± 11 | 1255 ± 9 | 1237 ± 7 | <**0.0001** | **0.0026** | 0.8485 |
| Ct.Por (%) | 0.435 ± 0.057 | 0.499 ± 0.075 | 0.446 ± 0.082 | 0.474 ± 0.48 | 0.0665 | 0.7656 | 0.4499 |
| pMOI (mm<sup>4</sup>) | 0.511 ± 0.070 | 0.484 ± 0.078 | 0.499 ± 0.077 | 0.511 ± 0.043 | 0.7678 | 0.7732 | 0.4419 |

Data presented as mean ± SD.
Ct. = cortical; Ar. = area; Tt. = total; Ma. = medullary; Ct.Ar/Tt.Ar= cortical area/total area; Ct.TMD = cortical tissue mineral density; Ct.Por. = cortical porosity; Ct.Th. = cortical thickness; pMOI = polar moment of inertia.
Two-way ANOVA: Bold values indicate significance with $p < 0.05$

Within the cortical bone of the femur, there was a significant main effect of housing condition across temperatures. Isolation decreased Ct.Ar., Ct.Ar./Tt.Ar., Ct.Th., and Ct.BMD, and increased Ma.Ar across temperatures. We also found a significant main effect of temperature in Ct.Th. and Ct.BMD, with thermoneutrality reducing Ct.Th. and Ct.BMD across housing conditions (see **Table 2**). There was no significant interaction between housing and temperature on cortical parameters.

Isolation and thermoneutrality had similar effects on the trabecular bone of the L5 vertebrae as in the femur (**Table 3**). Housing had a significant main effect in all L5 parameters except Tb.N. and Tb.Sp. There was a significant interaction between housing and temperature on BV/TV, BMD, BS/BV, SMI, and Tb.Th., with greater differences between isolated and grouped mice at room temperature. Isolation reduced BV/TV and BMD by approximately 22% and 21% at room temperature respectively, but only by approximately 10% for both parameters at thermoneutrality. Unlike the femur, however, there were no significant or trending differences between housing matched room temperature and thermoneutral mice. This suggests the significant interactions are driven both by a reduction in BV/TV and BMD in the grouped mice and an increase in these parameters in isolated mice at thermoneutrality.

**Table 3.**
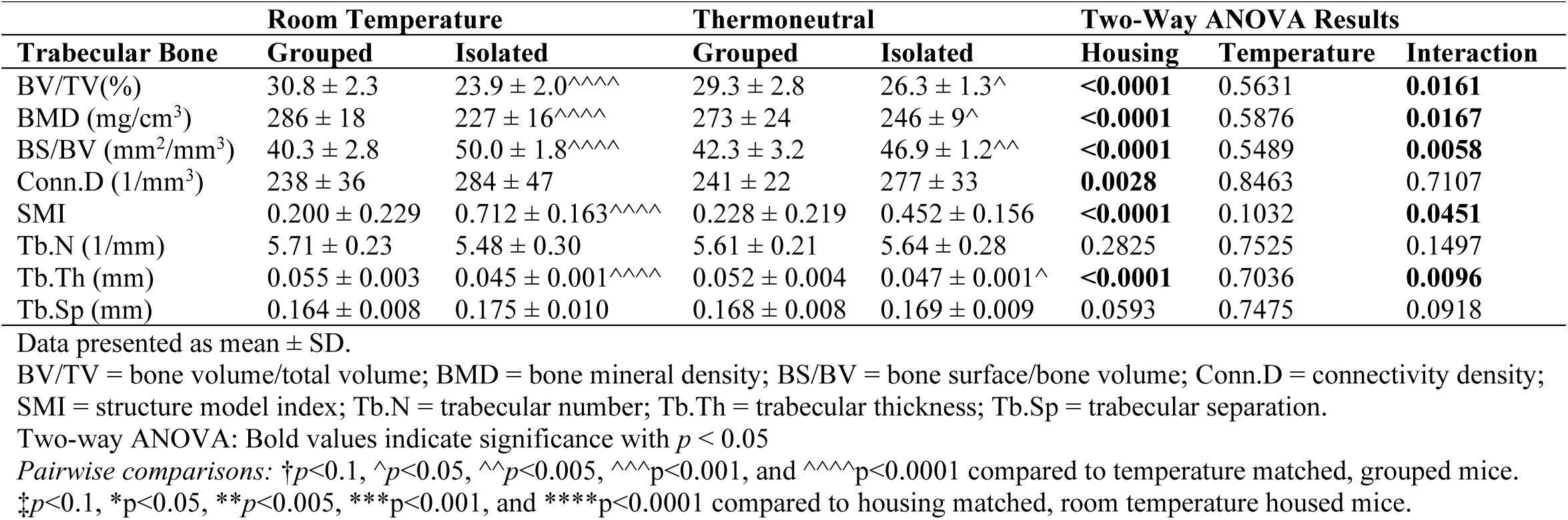
Trabecular microarchitecture of L5 vertebrae of male mice housed at either room temperature or thermoneutral in grouped or isolated housing for four weeks. (N=8/group).

There were no significant differences in weight or fat mass observed via DXA **(Supplementary Figure S1**). There was, however, a trending decrease in lean mass with thermoneutral housing (*p =* 0.0969).

### 3.2 Housing and temperature did not alter serum turnover markers

We examined bone turnover markers to determine if temperature or housing significantly altered bone formation or resorption markers. There were no significant main effects of either temperature or housing on P1NP or CTX-I (**Figure 2**). There were also no significant interactions between housing and temperature.

**Figure 2.**
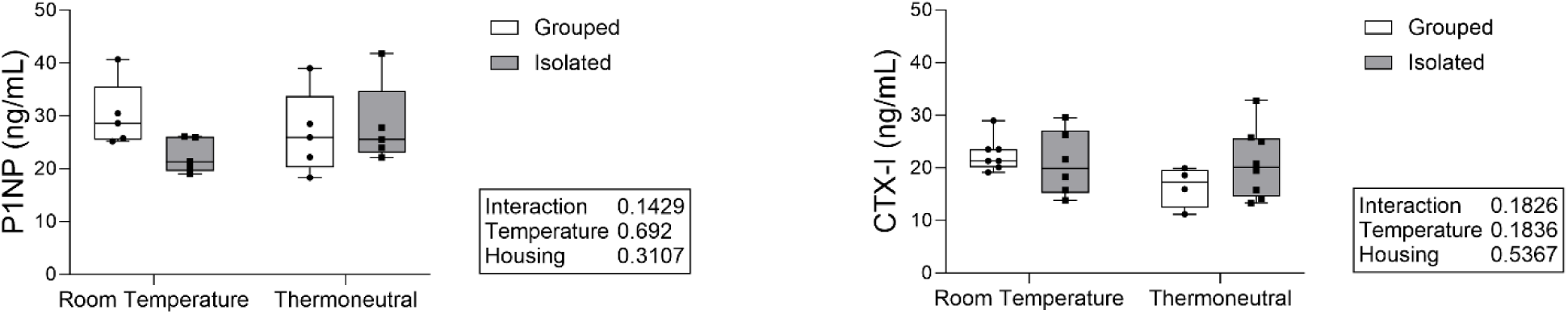
Social isolation and housing temperature did not significantly affect serum-level bone formation or resorption markers. Bone formation (P1NP) and resorption (CTX-I) markers were measured with serum EIA assays. N = 4-8/group.

### 3.3 Social isolation increased early osteoclast-related gene expression across housing temperatures

To further investigate changes in bone formation and resorption, we examined osteoblast and osteoclast-related gene expression in the whole tibia (**Figure 3**). Social isolation tended to increase expression of the early osteoclast-related gene NF-κB subunit RelA/p65 (*Rela*), and significantly increased expression of the M-CSF1 receptor (*Csfr1*), both important in osteoclast differentiation, independent of housing temperature. SH3 and PX domains 2A (*Sh3pxd2a*), important in osteoclast fusion and podosome formation, was significantly increased by isolation across housing temperatures. Social isolation also tended to decrease expression of osteoblast-related gene Runt-related transcription factor 2 (*Runx2*), important in osteoblast differentiation, independent of temperature. There was, however, no significant main effect or interaction of housing and temperature on RANKL (*Tnfs11*), or OPG (*Tnfrsf11b*). There were also no significant differences in expression of other osteoblast-related genes including *Bglap* or *Dmp1* (data not shown).

**Figure 3.**
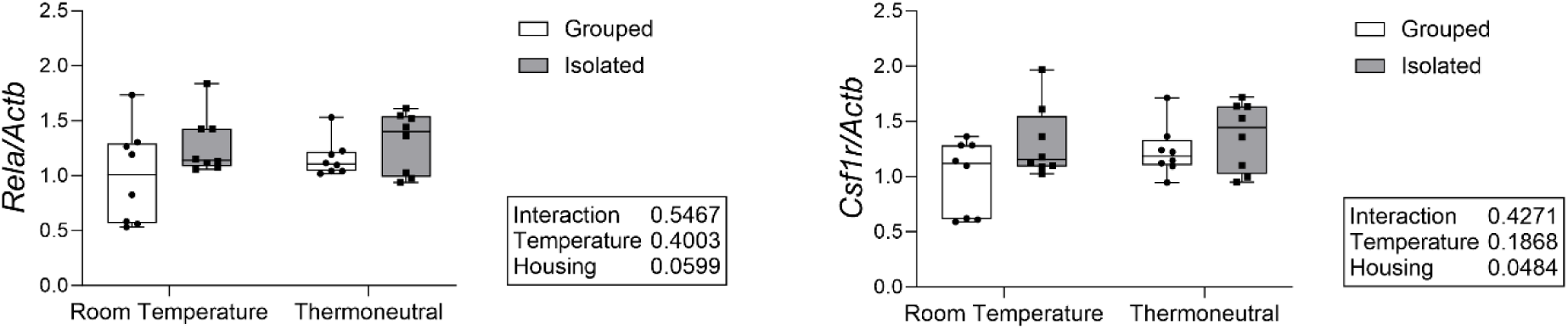
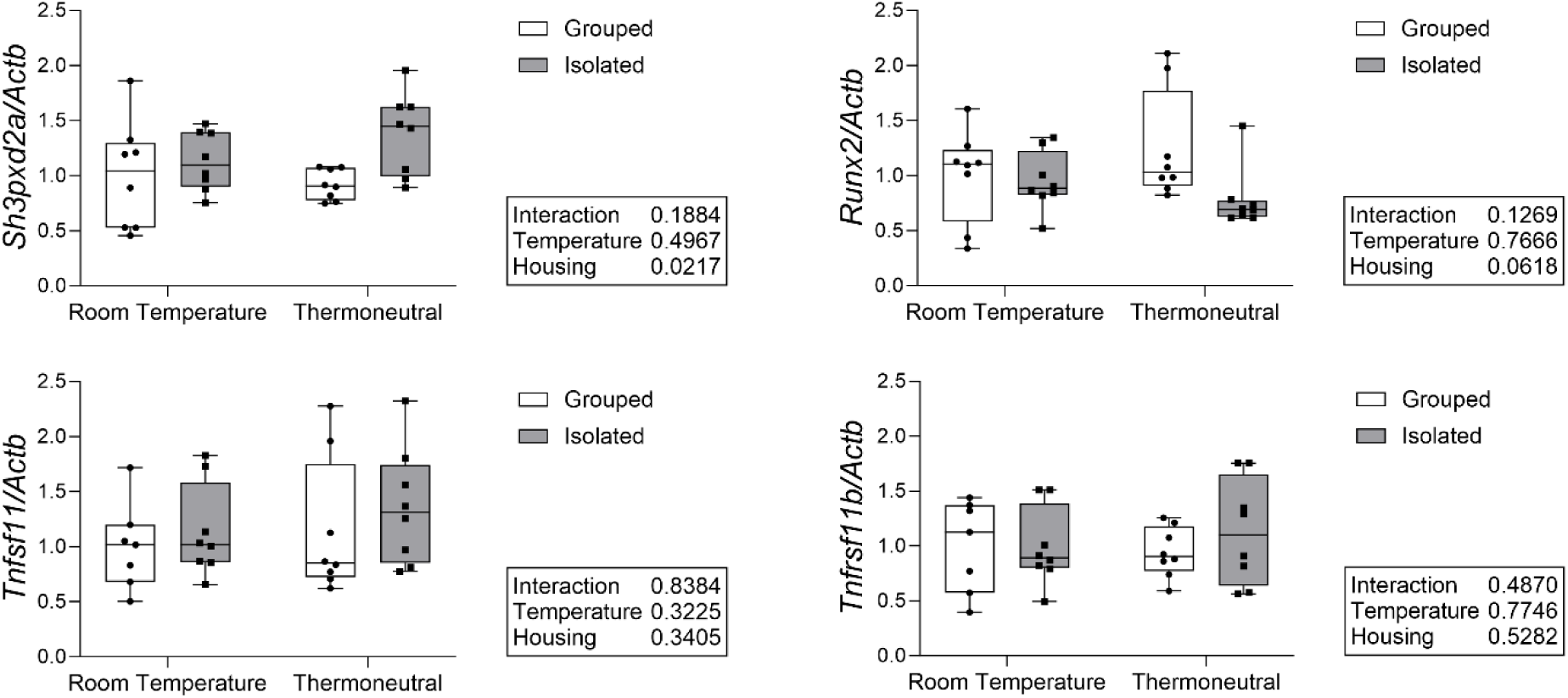
Social isolation increases early osteoclast related gene expression across housing temperatures. Measured in whole tibia with qPCR. Normalized to *Actb* as housekeeping gene. N=7-8/group.

### 3.4 Social isolation increased glucocorticoid-, but not sympathetic nervous system-related gene expression in bone across housing temperatures

To investigate whether the bone changes observed with isolation may be a result of glucocorticoid signaling, we measured expression of glucocorticoid-related genes in whole bone. Social isolation increased the expression of the glucocorticoid receptor (*Nr3c1*) independent of housing temperature (**Figure 4**). Isolation also tended to increase the expression of Hydroxysteroid 11-Beta Dehydrogenase 1 (*Hsd11b1*), an enzyme critical for corticosterone activation. The sympathetic nervous system is also an important component of the stress response and has significant effects on bone. We therefore also measured sympathetic nervous system-related gene expression, specifically the β1 and β2 adrenergic receptors (*Adrb1, Adrb2*), but found no significant effect of either housing or temperature.

**Figure 4.**
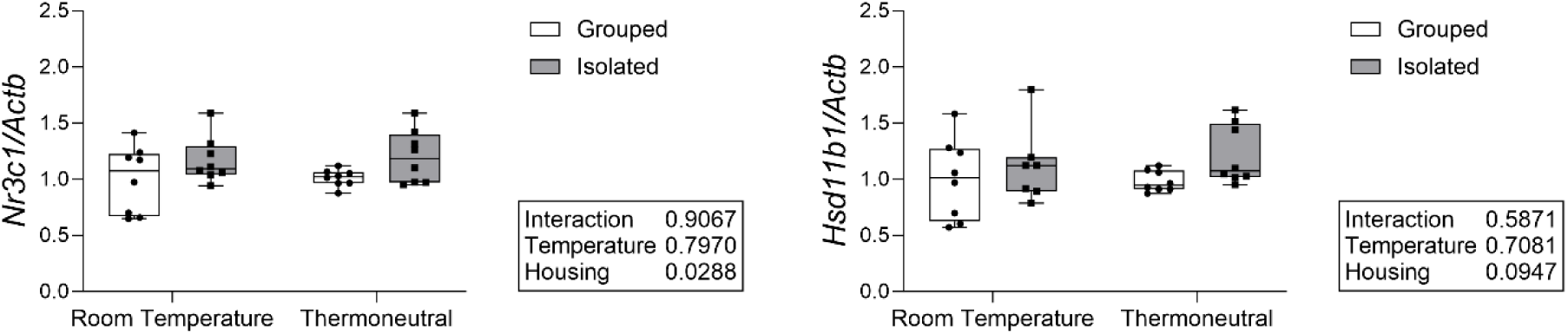
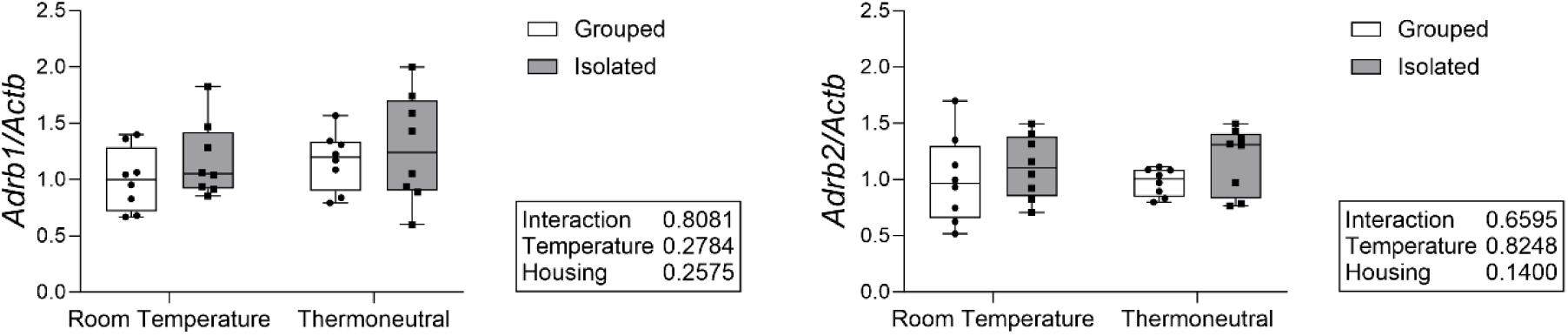
Social isolation increases glucocorticoid receptor expression across housing temperatures but does not affect sympathetic nervous system-related gene expression in whole bone. Measured in whole tibia with qPCR. Normalized to *Actb* as housekeeping gene. N=7-8/group.

### 3.5 Thermoneutral housing increased lipid area in BAT, while social isolation increased Pdk4 expression across housing temperatures

To ascertain if the observed changes in bone may be related to changes in adipose tissue, we compared BAT lipid density between groups. Thermoneutral housing increased lipid droplet percent area across housing groups, while isolation had no effect on percent lipid area (**Figure 5**).

**Figure 5.**
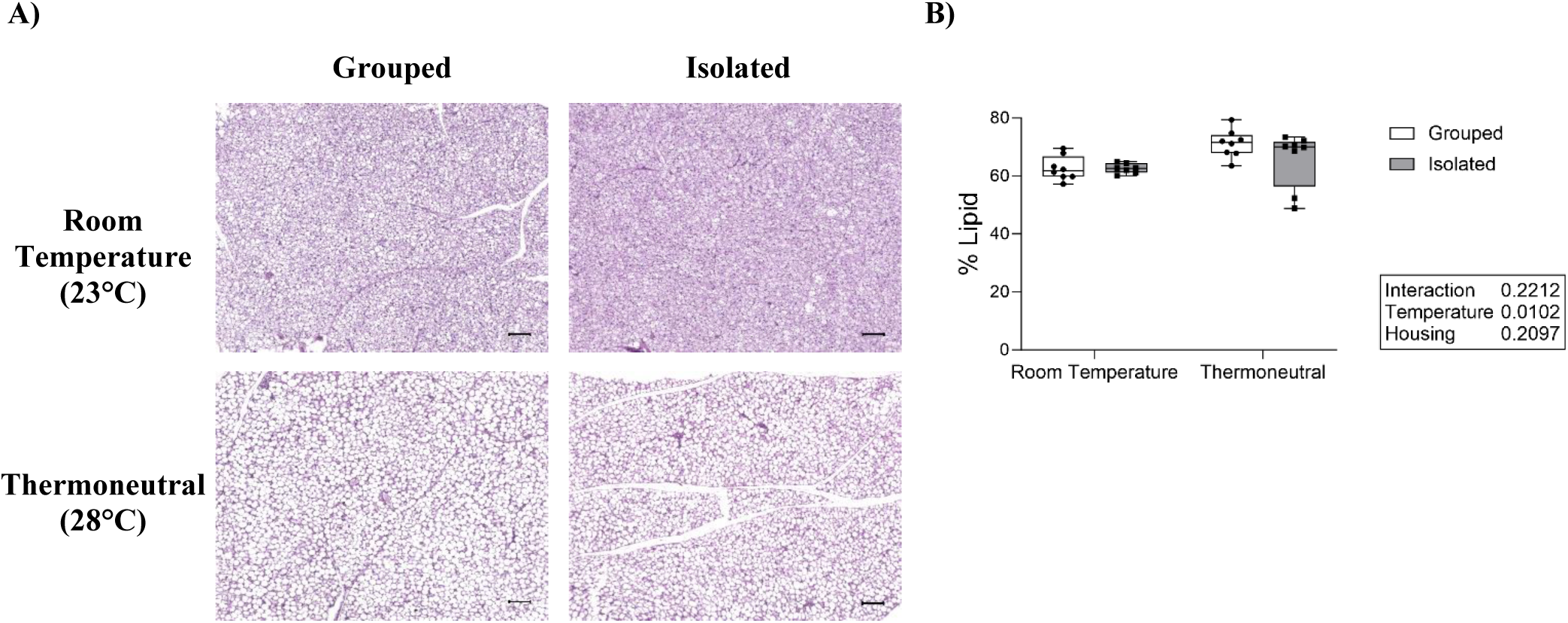
Thermoneutral housing increases percent lipid area. A) Representative images of H&E-stained brown adipose tissue (BAT). Imaged at 10x magnification, scale bar = 100µm. B) Percent lipid area of dissected BAT. N=8/group.

To further evaluate the effects of temperature and housing on BAT, we examined mitochondrial- and thermogenic-related gene expression in dissected BAT. Thermoneutral housing significantly decreased expression of type II deiodinase (*Dio2*), an enzyme that catalyzes T4 to T3 during sympathetic induced thermogenesis. Uncoupling protein 1 (*Ucp1*), which encodes a protein crucial for heat dissipation by H^+^ ion release across the inner mitochondrial membrane, was also significantly decreased by thermoneutral housing across housing conditions. Thermoneutrality also tended to decrease expression of pyruvate dehydrogenase lipoamide kinase isozyme 4 (*Pdk4*), which promotes oxidation of fatty acids upon lipolysis. (**Figure 6**). Social isolation, however, significantly increased *Ucp1* and *Pdk4* expression independent of housing temperature, suggesting social isolation may alter BAT metabolism independent of temperature. There was no significant effect of housing or temperature on expression of peroxisome proliferative activated receptor gamma coactivator 1-alpha (*Ppargc1a*), a transcriptional coactivator important in regulating genes involved in energy metabolism and thermogenesis.

**Figure 6.**
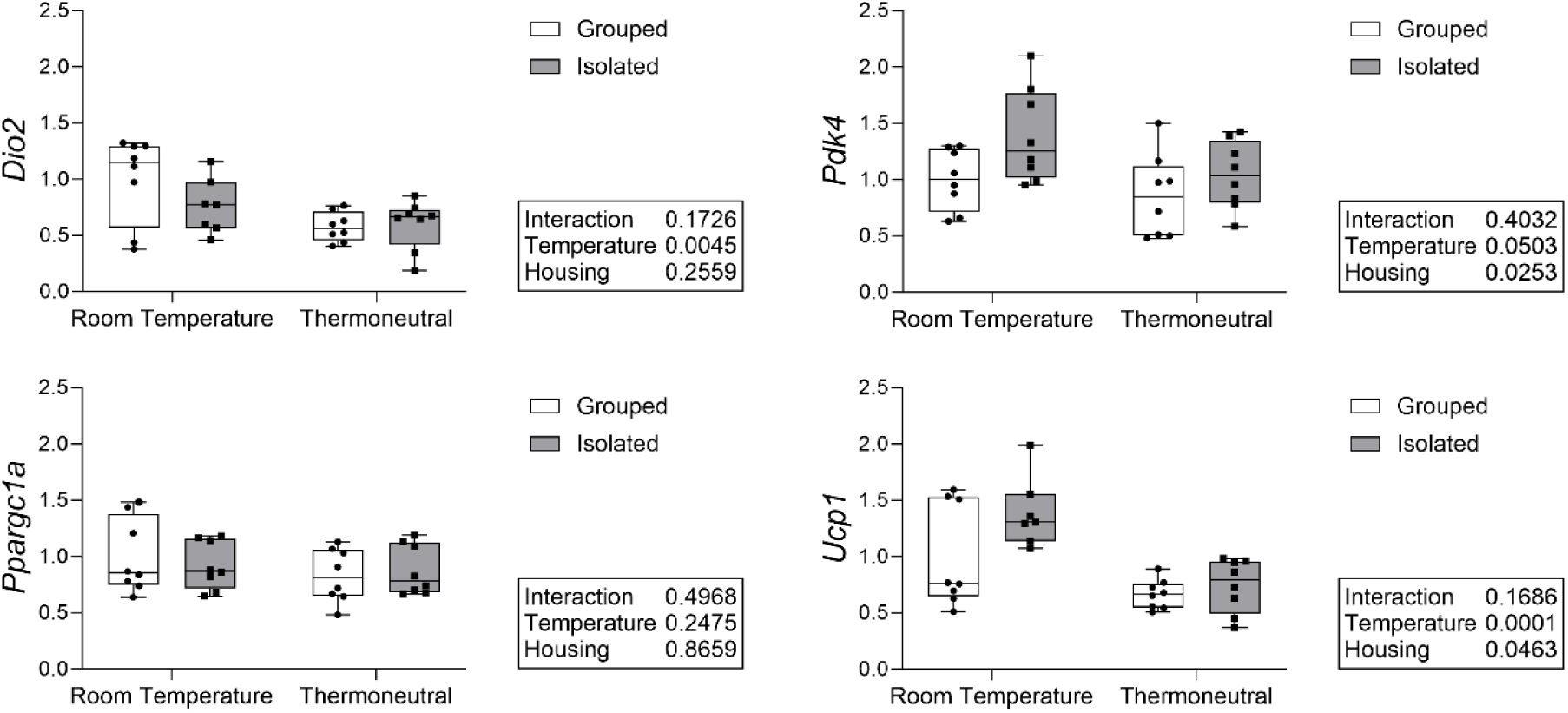
Thermoneutral housing decreased *Dio2*, *Pdk4*, and *Ucp1* expression, but isolation increased *Pdk4* expression across temperatures. Measured in brown adipose tissue (BAT) with qPCR. Normalized to highest CQ/lowest expressor. N=7-8/group.

## 4. Discussion

The major finding of this study is that thermoneutral housing does not fully rescue isolation-induced bone loss. Social isolation decreased trabecular and cortical parameters in the femur across temperatures and decreased trabecular parameters of the L5 vertebrae. There was a significant interaction effect between housing and temperature within some of the trabecular parameters, with a smaller effect of isolation at thermoneutrality. This may be suggestive of a partial amelioration of isolation-induced bone loss at thermoneutrality. However, in the femur, this interaction effect is driven largely by a trending reduction in bone parameters—specifically BV/TV and BMD—within the grouped-thermoneutral mice relative to the grouped-room temperature mice. This indicates the significant interaction between housing and temperature is driven both by a reduction in bone parameters in grouped mouse as well as an increase in isolated mouse bone parameters at thermoneutrality. While we did not see a significant or trending difference between the room temperature and thermoneutral grouped mice within the vertebrae, there was also no significant improvement within the isolated mice. This suggests that while thermoneutral housing may have some positive effects on bone parameters as previously demonstrated, these effects are limited within the isolated group and does not fully ameliorate the isolation-induced bone loss.

There were no significant main effects or interactions within the serum-level turnover markers.

We did find a significant increase in early osteoclast-related gene expression in the isolated mice independent of temperature. This may provide insight into the mechanisms underlying isolation-induced bone loss, but again suggests that thermoneutral housing does not fully ameliorate the effects of social isolation on bone.

We also examined gene expression related to glucocorticoid and sympathetic nervous system activity. We found a significant increase in glucocorticoid receptor expression and a trending increase in *Hsd11b1*, which is important in corticosterone activation, in the bone of isolated mice independent of temperature. We did not find any significant differences in expression of either of the beta-adrenergic receptors examined. This suggests that glucocorticoid signaling may play an important role in isolation-induced bone loss.

While thermoneutral housing did not fully rescue isolation-induced bone loss, within the BAT, thermoneutral housing increased percent lipid area, and decreased the expression of *Ucp1*. This is consistent with findings from previous thermoneutrality studies^(17,18)^ and suggests a reduction in cold stress at 28°C despite the lack of improvement in bone. We also found that social isolation increased expression of *Pdk4* across housing temperatures. While BAT activity and related thermogenesis are known to be mediated by the sympathetic nervous system^(19)^, glucocorticoid signaling can also increase BAT activity^(20)^, including the expression of *Pdk4*^(21)^. This further supports the hypothesis that glucocorticoid signaling may be an important mediation of social isolation-induced bone loss.

Although previous studies investigating the effects of thermoneutral housing on bone have found improvements in bone parameters, several studies have shown this is not always the case. Tastad et al. found that housing 10-week-old mice at 32°C for 6 weeks did not improve bone size or trabecular architecture within control groups or in tibial-loaded raloxifene treated mice^(22)^. In several cases, thermoneutral housing had a slight negative effect, including on cortical thickness, as seen in our own study. Our own work examining the effects of housing temperature on skeletal effects of the atypical antipsychotic olanzapine similarly found that thermoneutrality did not fully rescue olanzapine-induced bone loss^(23)^. In fact, olanzapine seemed to attenuate the positive effects of thermoneutral housing on bone.

Strengths of this study include that it is the first to examine the effects of temperature on social isolation-induced bone loss. Our results showed that bone loss persists at thermoneutrality and is likely not a result of thermal stress. We further showed evidence of elevated glucocorticoid receptor expression in isolated mice across temperatures, providing insight into possible mechanisms causing isolation-induced bone loss.

There were several limitations in this study that should be considered. First, it is possible that the thermoneutral housing was not sufficiently warm for the mice to achieve thermoneutrality. The thermoneutral range of mice has previously been defined as a range from approximately 26-34°C^(12)^.

However, Skop et al. suggests a diurnal 29-30°C is more accurate^(24)^, while other researchers house thermoneutral mice at ∼32°C^(17,25)^. We did observe changes in brown adipose tissue, including expansion of the lipid droplet size and decrease in *Ucp1* expression, consistent with thermoneutral temperature, although this did not rescue the effects of social isolation on bone. Furthermore, Sattgast et al. found that 26°C was sufficient to reduce trabecular bone loss^(18)^. It is also possible that housing mice in the warm temperature incubator induces additional stress relative to their prior environment within a larger room in the animal facility. If this is the case, this may mask the potential benefits of thermoneutral housing on isolation-induced bone loss.

A second limitation in this experiment was the use of only male mice. Our previous study found that social isolation through single housing only induced bone loss in male mice. We therefore only included males in this study to examine the potential ameliorating effects of thermoneutrality. We do not expect the results to be drastically different in females, as others have found no differences in the effects of temperature on bone between the sexes^(25)^. Some of our assays also had very low sample sizes, specifically our serum-level turnover markers had a N=<5 for some groups due to exclusion of coefficient of variation above 20, which may affect the results.

Future studies should consider the role of glucocorticoid signaling in isolation-induced bone loss, as well as the potential role of the sympathetic nervous system. Circulating measures of corticosterone and catecholamines, in addition to inhibition experiments, would help clarify the role of both glucocorticoid and sympathetic activity respectively. Additional experiments should also interrogate the sex differences in isolation-induced bone loss found in our previous study^(11)^. It is possible that social isolation acts on different time scales in female mice, potentially requiring longer periods of isolation to affect bone. Age may also play an important role, as social isolation at young ages, specifically post-weaning, has dramatic effects on brain development and behavior in rodents^(26)^, and may have similar effects on bone loss. Lastly, it is important to understand how isolation affects skeletal health in human populations. Only two studies have examined the effects of isolation on osteoporosis and fracture risk in humans^(9,27)^, and more work is needed particularly to understand the role of sex and age on these effects.

Overall, our results show that the negative effects of social isolation on bone persist even at thermoneutrality. This suggests that isolation-induced bone loss in mice is not a result of thermal stress due to single housing. Future work should focus on other potential mechanisms including the role of glucocorticoid signaling. Collectively, this study provides important insight into the mechanisms of social isolation-induced bone loss.

## Acknowledgments

The authors thank Grazina Armie Mangoba and Rea Anunciado-Koza for technical assistance.

## Data Availability

Data will be made available upon request.

## Supplementary Materials

**Supplemental Figure S1.**
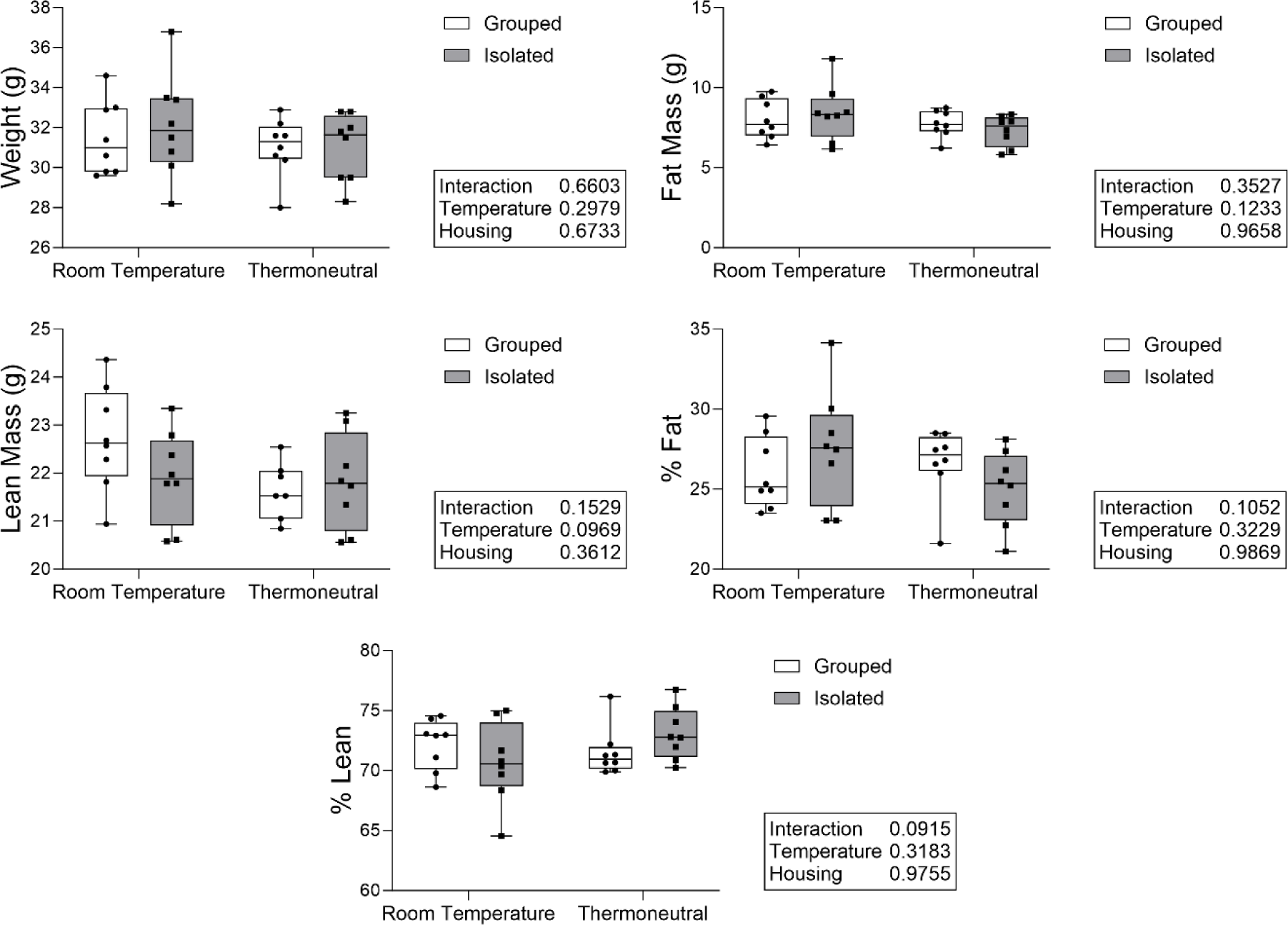
Housing and temperature did not significantly alter body parameters. At the end of 4 weeks of treatment, body parameters were measured using dual-energy x-ray absorptiometry (DXA). N=7-8/group.

**Supplementary Table S1.**
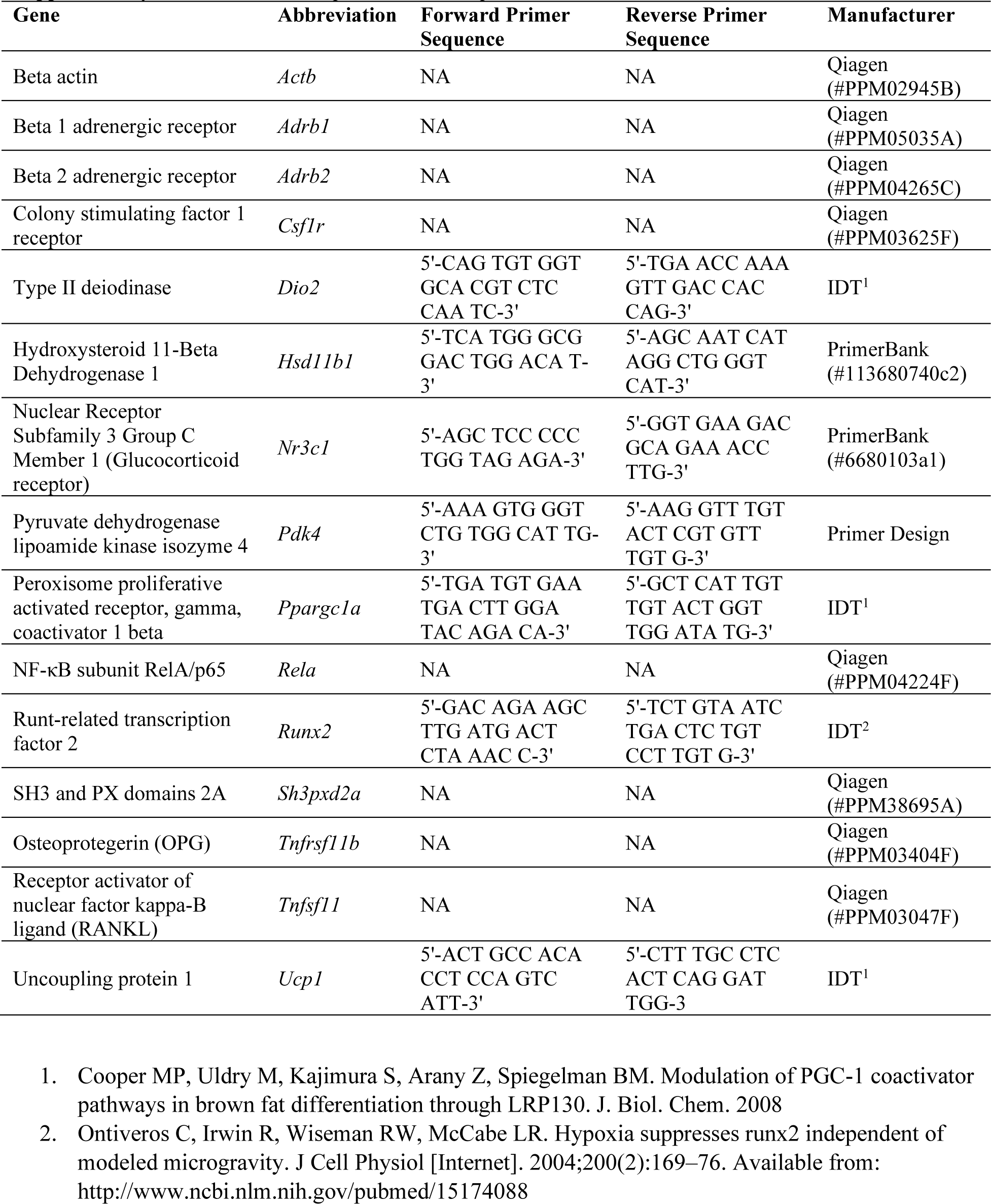
Primer sequences used for qPCR.

## References

1. Galambos C, Lubben J. Social Isolation and Loneliness in Older Adults: A National Academies of Sciences, Engineering, and Medicine Report. Innovation in Aging. 2020;4(Supplement_1):713-.

2. Kovacs B, Caplan N, Grob S, King M. Social networks and loneliness during the COVID-19 pandemic. Socius. 2021;7:2378023120985254.

3. U.S. Surgeon General. Our Epidemic of Loneliness and Isolation: The US Surgeon General’s Advisory on the Healing Effects of Social Connection and Community. Washington DC: US Department of Health and Human Services; 2023.

4. Valtorta NK, Kanaan M, Gilbody S, Ronzi S, Hanratty B. Loneliness and social isolation as risk factors for coronary heart disease and stroke: systematic review and meta-analysis of longitudinal observational studies. Heart. 2016;102(13):1009–16.

5. Holwerda TJ, Deeg DJ, Beekman AT, van Tilburg TG, Stek ML, Jonker C, et al. Feelings of loneliness, but not social isolation, predict dementia onset: results from the Amsterdam Study of the Elderly (AMSTEL). Journal of Neurology, Neurosurgery & Psychiatry. 2014;85(2):135–42.

6. Shen C, Rolls ET, Cheng W, Kang J, Dong G, Xie C, et al. Associations of social isolation and loneliness with later dementia. Neurology. 2022;99(2):e164–e75.

7. Donovan NJ, Wu Q, Rentz DM, Sperling RA, Marshall GA, Glymour MM. Loneliness, depression and cognitive function in older US adults. International journal of geriatric psychiatry. 2017;32(5):564–73.

8. Holt-Lunstad J, Robles TF, Sbarra DA. Advancing social connection as a public health priority in the United States. American psychologist. 2017;72(6):517.

9. Bevilacqua G, Jameson KA, Zhang J, Bloom I, Ward KA, Cooper C, et al. The association between social isolation and musculoskeletal health in older community-dwelling adults: findings from the Hertfordshire Cohort Study. Quality of life research. 2021;30(7):1913–24.

10. Schiavone S, Morgese MG, Mhillaj E, Bove M, De Giorgi A, Cantatore FP, et al. Chronic psychosocial stress impairs bone homeostasis: a study in the social isolation reared rat. Frontiers in Pharmacology. 2016;7:152.

11. Mountain RV, Langlais AL, Hu D, Baron R, Lary CW, Motyl KJ. Social isolation through single housing negatively affects trabecular and cortical bone in adult male, but not female, C57BL/6J mice. Bone. 2023;172:116762.

12. Gordon C. Thermal physiology of laboratory mice: defining thermoneutrality. Journal of thermal biology. 2012;37(8):654–85.

13. Du J, He Z, Xu M, Qu X, Cui J, Zhang S, et al. Brown Adipose tissue rescues bone loss Induced by Cold exposure. Frontiers in Endocrinology. 2022;12:778019.

14. Motyl KJ, Bishop KA, DeMambro VE, Bornstein SA, Le P, Kawai M, et al. Altered thermogenesis and impaired bone remodeling in Misty mice. Journal of Bone and Mineral Research. 2013;28(9):1885–97.

15. Appana B, Queen NJ, Cao L. Protocol to minimize the confounding effect of cold stress on socially isolated mice using thermoneutral housing. STAR protocols. 2023;4(3):102533.

16. Tero BW, Fortier B, Soucy AN, Paquette G, Liaw L. Quantification of lipid area within thermogenic mouse perivascular adipose tissue using standardized image analysis in FIJI. Journal of vascular research. 2022;59(1):43–9.

17. Iwaniec UT, Philbrick KA, Wong CP, Gordon JL, Kahler-Quesada AM, Olson DA, et al. Room temperature housing results in premature cancellous bone loss in growing female mice: implications for the mouse as a preclinical model for age-related bone loss. Osteoporosis International. 2016;27:3091–101.

18. Sattgast LH, Wong CP, Branscum AJ, Olson DA, Aguirre-Burk AM, Iwaniec UT, et al. Small changes in thermoregulation influence cancellous bone turnover balance in distal femur metaphysis in growing female mice. Bone Reports. 2023;18:101675.

19. Collins S. β-Adrenergic receptors and adipose tissue metabolism: evolution of an old story. Annual review of physiology. 2022;84:1–16.

20. Ramage LE, Akyol M, Fletcher AM, Forsythe J, Nixon M, Carter RN, et al. Glucocorticoids acutely increase brown adipose tissue activity in humans, revealing species-specific differences in UCP-1 regulation. Cell metabolism. 2016;24(1):130–41.

21. Connaughton S, Chowdhury F, Attia RR, Song S, Zhang Y, Elam MB, et al. Regulation of pyruvate dehydrogenase kinase isoform 4 (PDK4) gene expression by glucocorticoids and insulin. Molecular and cellular endocrinology. 2010;315(1-2):159–67.

22. Tastad CA, Kohler R, Wallace JM. Limited impacts of thermoneutral housing on bone morphology and mechanical properties in growing female mice exposed to external loading and raloxifene treatment. Bone. 2021;146:115889.

23. Langlais AL, Mountain RV, Kunst RF, Barlow D, Houseknecht KL, Motyl KJ. Thermoneutral housing does not rescue olanzapine-induced trabecular bone loss in C57BL/6J female mice. Biochimie. 2023;210:50–60.

24. Škop V, Guo J, Liu N, Xiao C, Hall KD, Gavrilova O, et al. Mouse thermoregulation: introducing the concept of the thermoneutral point. Cell reports. 2020;31(2).

25. Martin SA, Philbrick KA, Wong CP, Olson DA, Branscum AJ, Jump DB, et al. Thermoneutral housing attenuates premature cancellous bone loss in male C57BL/6J mice. Endocrine Connections. 2019;8(11):1455–67.

26. Fone KC, Porkess MV. Behavioural and neurochemical effects of post-weaning social isolation in rodents—relevance to developmental neuropsychiatric disorders. Neuroscience & Biobehavioral Reviews. 2008;32(6):1087–102.

27. Lee A, McArthur C, Ioannidis G, Mayhew A, Adachi JD, Griffith LE, et al. Associations between Social Isolation Index and changes in grip strength, gait speed, bone mineral density (BMD), and self-reported incident fractures among older adults: Results from the Canadian Longitudinal Study on Aging (CLSA). PLoS one. 2023;18(10):e0292788.

